# Localising elements in single-particle reconstructions by REEL-EM: Reconstructed Electron Energy-Loss - Elemental Mapping

**DOI:** 10.1101/2024.01.18.575858

**Authors:** Olivia Pfeil-Gardiner, Higor Vinícius Dias Rosa, Dietmar Riedel, Yu Seby Chen, Dominique Lörks, Pirmin Kükelhan, Martin Linck, Heiko Müller, Filip Van Petegem, Bonnie J. Murphy

## Abstract

For structures determined by single particle cryo-EM, no technique currently exists for mapping elements to defined locations, leading to errors in the assignment of metals and other ions, cofactors, substrates, inhibitors, and lipids that play essential roles in activity and regulation. Elemental mapping in the electron microscope is well established for dose-tolerant samples but is challenging for biological samples, especially in a cryo-preserved state. Here, we combine electron energy-loss spectroscopy (EELS) with single-particle image processing to allow elemental mapping in cryo-preserved macromolecular complexes. Proof-of-principle data show that our method, REEL-EM, allows 3D reconstruction of EELS data, such that a high total electron dose is accumulated across many copies of a complex. Working with two test samples, we demonstrate that we can reliably localise abundant elements. We discuss the current limitations of the method and potential future developments.

## Introduction

In cryogenic electron microscopy (cryo-EM) single-particle analysis (SPA), experimental densities are interpreted to atomic models with the help of substantial prior information. This comprises knowledge of peptide or nucleotide sequence, bond lengths and angles, and secondary structure. For protein and nucleotide scaffolds, individual atoms can thus be identified, even though no tools exist for elemental mapping, and these atoms can be accurately placed, even though experimental data seldom reach atomic resolution. For all other components of a structure, including bound cofactors, lipids, substrates and inhibitors, posttranslational modifications, and especially metals and other ions, prior information is less easily available. For these, the interpretation of experimental data is thus often ambiguous, leading to errors in atomic models.

A recent study, analysing 30 randomly selected metalloproteins whose X-ray crystal structures had been deposited in the Protein Data Bank (PDB), found that a third to one half of the structures contain errors in metal assignment [1]. For structures determined by SPA cryo-EM, which in 2023 encompassed over a third of all deposited structures [2], the error rate is likely higher, since there is currently no method that retrieves elemental information in the spatial context of the structure – a challenge we aim to address.

Methods for bulk elemental analysis of purified complexes (X-Ray fluorescence or absorbance, optical emission spectroscopy, inductively coupled plasma mass spectrometry) cannot distinguish between specifically bound and soluble components and are inadequate for complexes with multiple binding sites. For structures determined by X-ray crystallography, the anomalous scattering signal at specific wavelengths offers a solution for many elements, though not for light elements (like C, N, O, Na, Mg, P) that are most abundant in biological samples. However, not all samples are amenable to crystallisation.

In the electron microscope, elemental information is available from inelastic scattering processes by means of energy-dispersive X-ray spectroscopy (EDX) and electron energy-loss spectroscopy (EELS) – both of which are recorded in scanning mode – or the corresponding imaging mode, electron spectroscopic imaging (ESI). These techniques have rarely been used for radiation-sensitive biological samples, as very high doses are required to accumulate sufficient signal from inelastic scattering events. A study evaluating the sensitivity of EELS for biological samples estimated that 10^7^ e^-^/Å^2^ would be required for single-atom detection [3]. Such a dose would leave little to no structural context intact to be analysed.

In cryo-preserved cell and tissue samples, abundant elements have been mapped by EDX, EELS, and ESI [4-12]. In one instance, the assignment of an unknown cofactor in a cryo-preserved protein sample was achieved on the basis of bulk EELS without any spatial information [13]. Only a few studies have evaluated three-dimensional (3D) elemental mapping by EM in macromolecular complexes [14-16].

In SPA, two-dimensional (2D) images with low signal-to-noise ratio (SNR) are combined into a single 3D reconstruction with high SNR. A similar reconstruction technique could conceivably be applied to accumulate spatially resolved elemental signals. Early studies of freeze-dried ribosomes and nucleosomes applied SPA to ESI images at the phosphorus edge to map nucleic acids in these complexes [14, 15]. However, the high doses necessary to estimate particle poses from ESI images are incompatible with cryo-preserved samples or high resolution. A separate study demonstrated the localisation of metal ions within a protein reconstruction exploiting the differences of light and heavy elements in high-angle scattering intensities, but did not discriminate between different heavy elements in a single reconstruction [16].

We propose a method that we call Reconstructed Electron Energy-Loss Elemental Mapping (REEL-EM) for a holistic elemental description of a structure in 3D. In this approach, we combine scanning transmission electron microscopy (STEM-) EELS with an SPA-like workflow to meet the high dose requirements for elemental mapping. A key difference to previous studies is that SPA particle poses are estimated on the basis of reference images formed by elastic scattering rather than from the energy-loss images themselves. Using these poses, 3D reconstructions can be generated for the full energy-loss spectrum, allowing analysis of elemental signatures in a 3D spatial context (Figure 1) at doses compatible with cryopreserved biological samples. To usefully address many biological questions, the technique would need to reach single-atom sensitivity. Although high resolution would also be desirable, even at low resolution, elemental signal can be overlaid on higher-resolution densities from standard SPA to inform the process of atomic model building. Therefore, the required resolution depends strongly on the individual biological question to be addressed.

**Figure 1.**
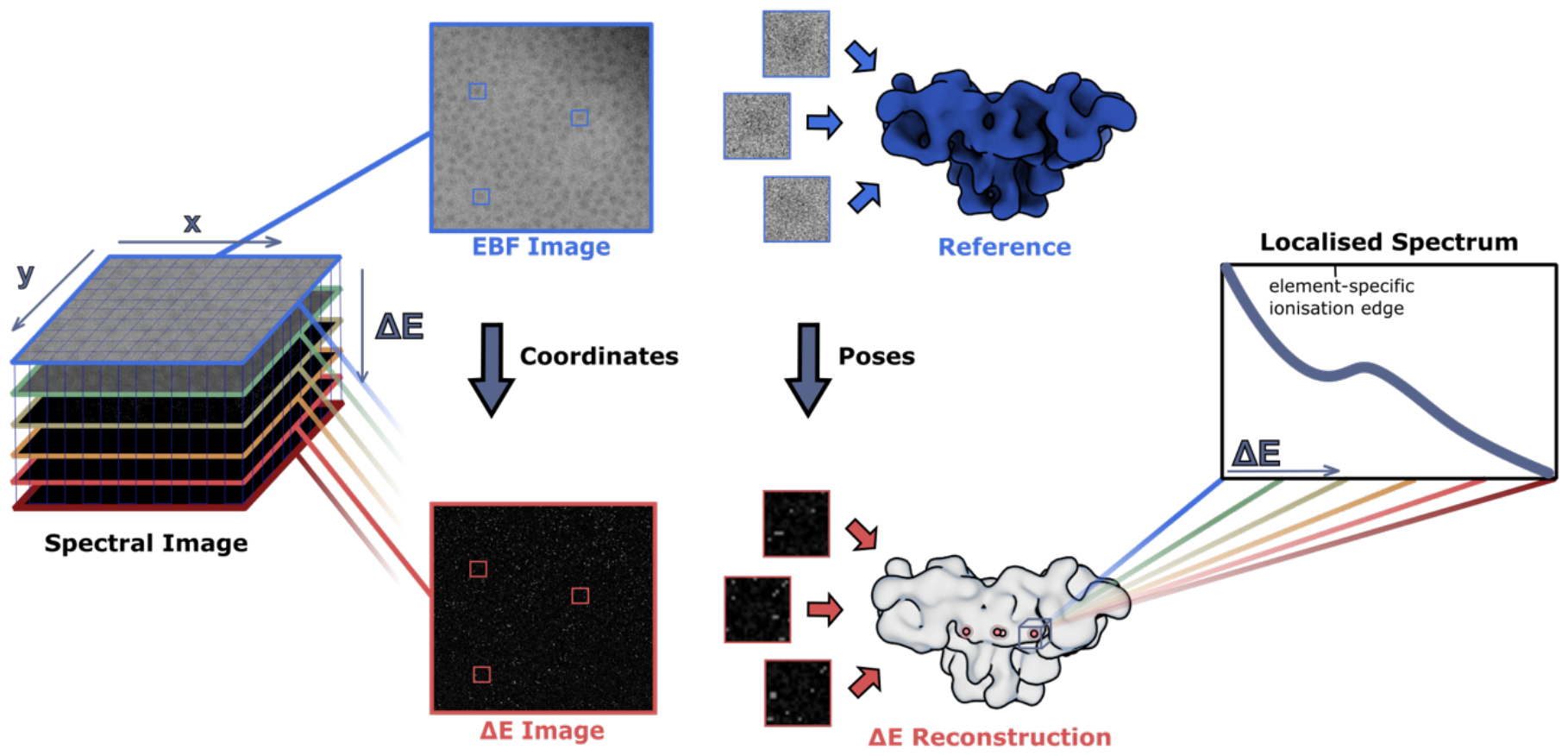
Schematic of the REEL-EM workflow. Spectral images of cryopreserved macromolecular complexes are collected in STEM-EELS mode. In addition to two spatial dimensions (x and y), these contain an energy-loss dimension (ΔE). From these images, the zero-loss portion is extracted to form elastic bright field images (EBF) from which particle positions and reconstruction poses can be determined by refining a reference reconstruction. Then, ΔE images are extracted for all energy bins along the energy-loss spectrum. Particle coordinates and reconstruction poses determined from the reference are then applied to each of these images – indicated here at one exemplary energy loss – such that a volume can be reconstructed for every energy loss, generating a 4D data set with three spatial dimensions and one energy-loss dimension. In addition to analysis of the reconstructions, localised spectra can be generated by plotting the values of specific voxels or voxel regions as a function of energy loss.

Here, we establish a workflow for data collection and processing for REEL-EM. Using this workflow, we provide proof-of-principle data demonstrating the reconstruction of specific elemental signals in the 3D particle space. We investigate the potential and current limitations of the method.

## Results

In order to enable elemental mapping of dose-sensitive complexes by REEL-EM, we have established the following workflow (Figure 1). We collect spectral images of a conventionally prepared single-particle sample, operating the microscope in STEM-EELS mode at doses below 100 e^-^/Å^2^. We sum intensities of the zero-loss portion of the spectral images to form high-intensity reference images that we use to determine particle coordinates and reconstruction poses. These coordinates and poses are subsequently used to reconstruct 3D volumes at each energy bin across the full energy-loss spectrum. The result is a four-dimensional (4D) data set containing three spatial and one energy-loss dimension(s). This can be viewed as a series of reconstructions, or as one reconstruction with a spectrum at each voxel. By analysing reconstructions at different energies or spectra at different locations, elements can be identified and mapped to spatial regions of a protein.

### A novel hardware configuration for low-noise, high-speed collection of cryo-STEM-EELS data

We carried out this work on a high-end electron microscope with a cryogenic stage. Relative to a microscope used for conventional SPA, several hardware and software modifications are required to collect low-noise spectral images. EELS spectra have to be collected pixel-by-pixel in STEM mode, requiring the respective alignments and scanning functionality. We found an annular dark field detector useful for faster navigation and focusing. An energy spectrometer with an energy resolution of at least 1eV and stability over several days is desirable. The detector should fulfil several requirements: The readout noise should be low to allow accurate detection of single electron events. This is especially important for the very low-dose acquisitions used in this method, where there are only a few electron events in the core-loss region of a given spectrum. The dynamic range should be large so that the intense zero-loss peak can be recorded simultaneously. Additionally, high frame rates are required for reasonable acquisition speeds as a detector readout is required at every scanning position and, therefore, 16 million times per 4kx4k spectral image, making the acquisitions slow. All these requirements are met by modern hybrid pixel detectors. We used a Titan Krios G3 in STEM mode, equipped with conventional annular dark field detectors, a CEOS CEFID energy filter [17] and a post-filter Dectris ELA hybrid pixel detector [18] for spectral image acquisition. In an attempt to balance detector performance with aberration effects, we chose an acceleration voltage of 200 kV for our experiments.

### Establishing an automated workflow for data collection

REEL-EM depends on large data sets to accumulate sufficient signal for elemental detection, which is substantially facilitated by automated acquisition of spectral images (see Figure S1). For this, we use a combination of the microscope automation software SerialEM and the filter-control software Panta Rhei. SerialEM is used for setting up and navigating acquisition positions and for automatically focusing on a carbon area adjacent to the acquisition position. SerialEM then calls an external python script that triggers spectral image acquisition and saving in Panta Rhei via its RPC (remote procedure call) interface. After the acquisition, the stage is moved to the next position and autofocus is carried out by SerialEM while, in parallel, Panta Rhei is assembling and saving the previous spectral image. In this way, the acquisition can run for several days, producing one 4kx4k spectral image every 23 minutes. The speed of the data collection is limited by the pixel dwell time, which depends on the maximum frame rate of the detector, and by the assembly and saving time for a spectral image in Panta Rhei. We acquired the spectral images with a dwell time of 30 µs (∼33 kHz), which is beyond the specification for the Dectris ELA, but we did not observe data loss in this mode. We found that the addition of gold fiducials to the carbon film aids the autofocus routine, which is, in our experience, much less robust in STEM than in TEM.

### Determining particle poses on the basis of elastic bright-field reference images

To calculate particle poses, a reference image is required, which could be derived in a number of ways. A conventional STEM detector records amplitude contrast signals due to differences in scattering angle, with elastic scattering angles being typically higher than inelastic scattering angles. The signal of elastically scattered electrons can be collected either by recording their presence at higher scattering angle, forming an annular dark field (ADF) image, or their absence at lower angles, forming a bright-field image. In our experiments, the camera length was adjusted so that the full bright-field disk entered the entrance aperture of the energy filter, excluding (mostly elastically-scattered) electrons scattered at higher angles (convergence angle α = 6 mrad; collection angle β = 8 mrad). Summing the counts in the zero-loss portion of the spectral image therefore gives an elastic bright field (EBF) image.

It is possible to collect an ADF image in parallel to spectral image acquisition. For a small data set of 3,086 particles of worm haemoglobin (erythrocruorin) from *Lumbricus terrestris*, we compared the quality of reconstructions, aligned with a Class3D job, from simultaneously collected dark-field and bright-field images. The collection angle range for the dark field detector was 15 – 150 mrad. We also compared a reconstruction from images for which the ADF images were subtracted from the EBF images (Figure 2). The bright-field images performed best, resolving most features in the reconstruction. The dark-field and mixed reconstructions, in comparison, appear more noise-dominated. This is most likely a consequence of larger amounts of detector noise in the ADF than in the EBF images. For the larger data sets, we therefore did not acquire simultaneous dark-field images and used only the EBF images as reference.

**Figure 2.**
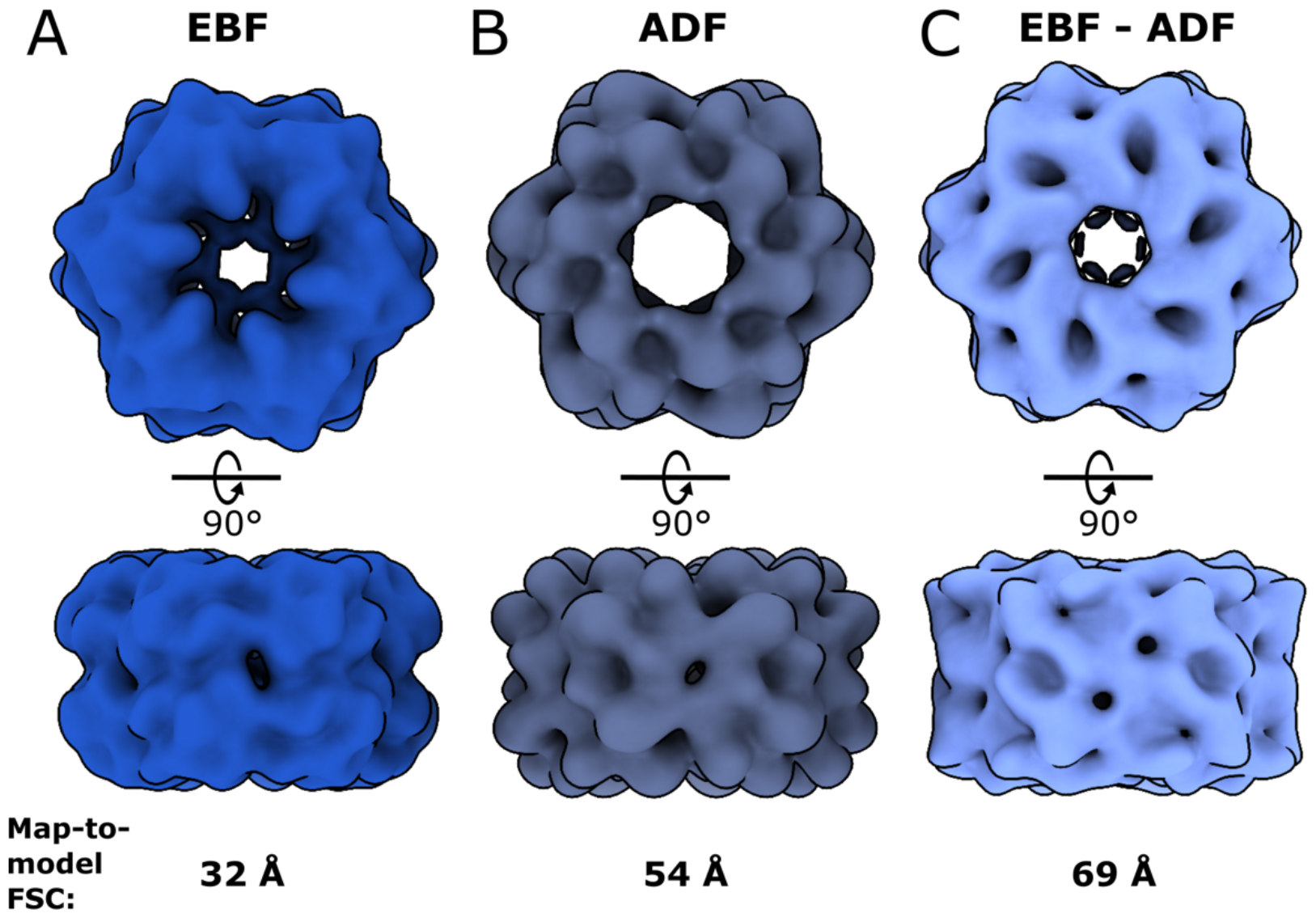
Evaluating the use of different image modalities for the estimation of reconstruction poses. SPA reconstructions of worm haemoglobin were refined from the same subset of 3086 particles which had been extracted from images collected simultaneously, but at different detectors. (A) The reconstruction refined from elastic bright field images, which were extracted from the zero-loss region of spectral images. (B) The reconstruction refined from annular dark-field images. (C) The reconstruction refined from combined images, generated by subtracting the annular dark-field images from the elastic bright-field images. The volume shown in A resolves the most features of worm haemoglobin, showing a better-quality reconstruction than B or C and a higher-resolution map-to-model FSC (PDB-5M3L) [34, 35] at a 0.5 cut-off.

From a data set of 1,142 spectral images of rabbit Ryanodine Receptor 1 (RyR1), we extracted EBF images from an energy region of ±3.6 eV. We picked 460,651 particles with a manually trained model in crYOLO [19], which we extracted and subjected to 2D classification for cleaning, 3D classification for pre-alignment and refinement in RELION [20], using an initial model generated from the data, low-pass filtered to 60 Å, as reference. No contrast transfer function (CTF) correction was applied during processing. The refinement yielded a reconstruction at 24 Å resolution after post-processing (Figure 3). The same procedure was performed on a smaller data set of worm haemoglobin (WH). For this sample, 16,648 EBF particle images yielded a resolution of 27 Å after post-processing (Figure S2).These refinements provided particle positions and poses to allow reconstruction of energy-loss data.

**Figure 3.**
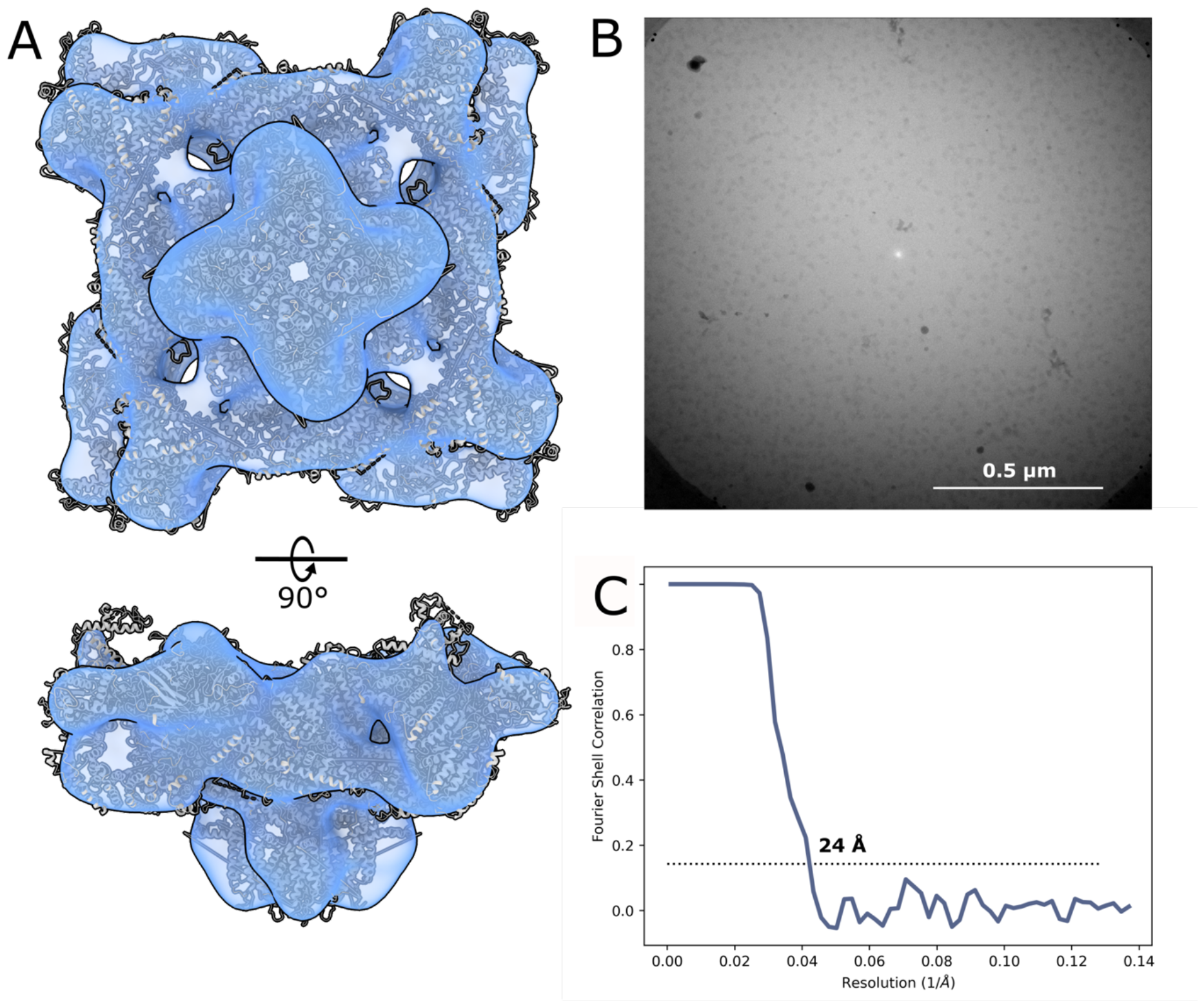
Reference reconstruction of RyR1. (A) Refined reconstruction of RyR1 from a data set of 460,651 particle images which were extracted from the elastic bright field image generated from zero-loss peak counts. The map shows a good fit - albeit at low resolution - to the published model of rabbit RyR1, PDB-5TAQ [22, 38], which is displayed as cartoon model. The refined poses can be used for reconstruction of energy-loss volumes. (B) An exemplary extracted EBF micrograph. (C) The gold-standard Fourier shell correlation for this reconstruction has a 0.143 cut-off at 24 Å.

### Reconstruction of energy-loss data generates 3D EELS maps

A schematic of the reconstruction procedure is shown in Figure S3. Star files from the reference refinements were manipulated to point to energy-loss images for each 0.77 eV energy bin of the spectral images. For each energy bin, particles were extracted and reconstructed to a 3D volume, using the command relion_reconstruct. To maintain interpretable EELS spectra after reconstruction, the volumes should only undergo linear transforms so that an external intensity scale is maintained. These terms are not met by relion_reconstruct, as this function sets the Fourier origin voxel to zero, thereby normalising the intensity of the reconstruction. We compiled a version of RELION without this normalisation and used this version for reconstruction. This modification comes at a cost of artefactual density, which is strongest at the corners of the reconstructed box (Figure S4).This effect can be minimised by oversampling the Fourier volume using the pad option. We found that a padding factor of 8 renders the intensity of the artefact sufficiently low for it not to be visible at threshold levels used for inspecting the reconstructions, although this resulted in noisier spectra. For this reason, all spectra displayed in this work are from reconstructions performed with 2-fold zero-padding and all displayed volumes from reconstructions performed with 8-fold zero-padding for better visualisation. The modified reconstruction procedure maintains the intensity profile of the EELS spectrum of the constituent particle images (Figure 4).

**Figure 4.**
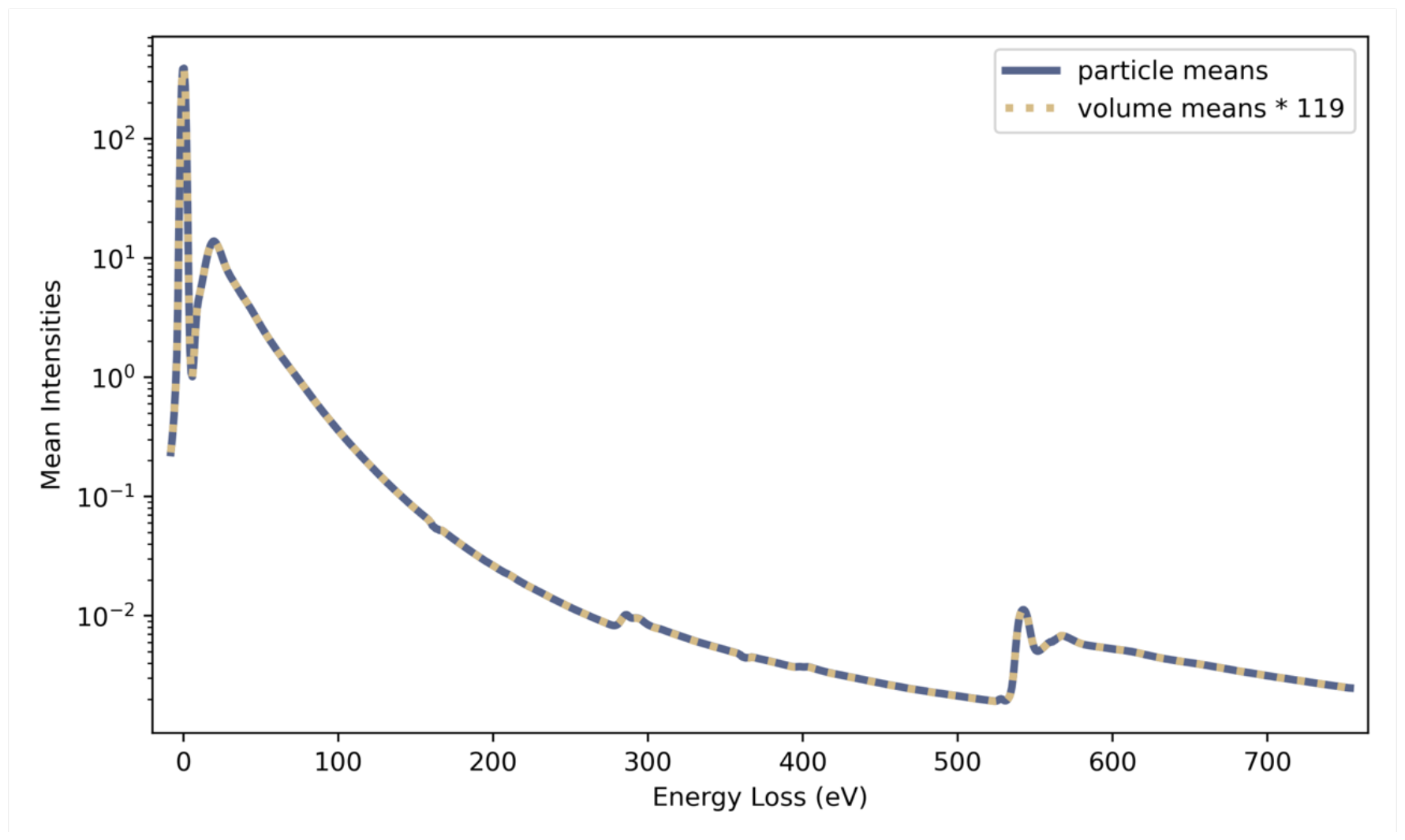
The mean intensities of particle images and respective 3D reconstructions maintain the same profile along the energy-loss spectrum. Mean intensities of all voxels in all particle images at each energy loss are indicated with the continuous blue line and mean intensities of the volumes, reconstructed from these images with a modified version of relion_reconstruct, are indicated with the green dashed line. Applying an empirical constant scaling factor of 119 to the mean intensities of the reconstructions, the curves become nearly identical, showing that the EELS profile is maintained by the reconstruction procedure.

Analysing the spectrum summed across all voxels of the RyR1 reconstructions, ionisation edges for the most abundant elements are clearly visible: The K-edges of carbon at 284 eV, oxygen at 532 eV (with a small pre-peak indicating the expected presence of gaseous oxygen due to radiation damage [21]), and nitrogen at 401 eV, as well as the L2,3-edges of the buffer components chlorine at 200 eV, potassium at 294 eV, and phosphorus at 132 eV (Figure 5). Movie 1 shows a sequence of all energy-loss reconstructions. There are evident changes in the 3D localisation of the signal with changing energy loss. These changes coincide with the major features in the spectrum: In the core-loss region prior to the carbon edge (237.5 – 242.1 eV), the intensity shows no apparent correlation with the complex. In contrast, at the carbon edge (283.2 – 287.8 eV), the density colocalises with the protein and the micelle. At the oxygen edge (540.4 – 545.0 eV), the density shows an inverse correlation with the complex, in line with the higher oxygen concentration in the solvent (Figure 5). These differences reflect the known elemental distributions of the complex. In addition to elemental features, the low-loss region (ΔE ≤ 50 eV) of the spectrum contains specific plasmon absorbances that are spatially resolved in the reconstructions (Figure S5).

**Figure 5.**
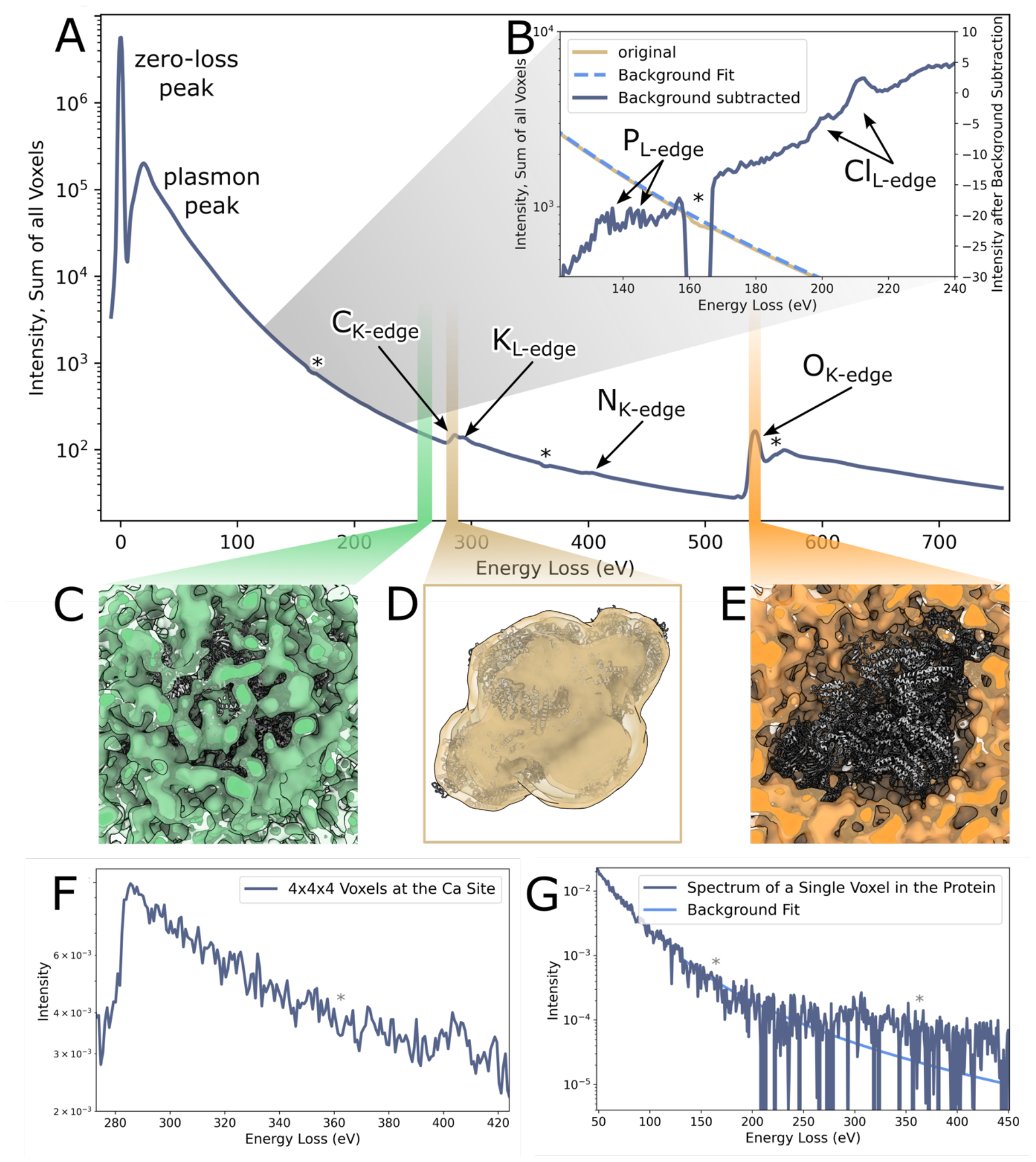
Selected spectra and volumes of reconstructed EELS data of RyR1. (A) The spectrum of the sum of all intensities in the EELS reconstructions shows elemental edges for the most abundant elements: carbon, potassium, nitrogen, and oxygen. The asterisks mark regions where the spectra show dips due to double-wide pixels at the tile boundaries of the detector, which are not perfectly accounted for by the detector’s flatfield correction. (B) For the region between 121 and 240 eV a background subtraction was performed, based on a fit of the spectrum between 121 eV and 127 eV. The subtracted spectrum shows additional edges for phosphorus and chlorine. (C-E) Gaussian-filtered reconstructions for three 4.6 eV-regions of the spectrum are shown together with an atomic model of rabbit RyR1, PDB-5TAQ [22, 38]. Before the carbon edge (C), the reconstruction shows a uniform distribution of intensity between the protein and the solvent. At the carbon edge (D), the reconstruction shows colocalization with the protein and micelle, and at the oxygen edge (E), the reconstruction shows colocalization with the solvent. The reconstructions reflect the known local distributions of carbon and oxygen. (F) The spectrum of summed intensities of a 4×4×4 voxel region at the known calcium-binding site does not show a calcium edge discernible above noise at 346, 350 eV. (G) The spectrum of a single voxel within the protein shows a clearly discernible carbon edge. A background fit based on the region 120 – 240 eV is displayed for better visualisation.

In principle, EELS analysis can detect all elements other than hydrogen (except as H2). Of the known elements in our sample we detect all except hydrogen, zinc, sulphur, and calcium. As the energy range and spectrum positioning on the detector were optimised for other edges in this data set, we do not expect to detect zinc because its major absorption edge (1020 eV, 1043 eV) lies beyond our chosen energy range, or sulphur (165 eV) because the associated energy loss coincides with an artefact at the detector tile edge (see for example Figure 5).The spectrum of a 4×4×4-voxel region near the known calcium-binding site formed by residues including E3893, E3967, and T5001 [22] does not show a calcium L2,3-edge signal discernible above noise, indicating that we have not reached single-atom sensitivity with this data set (Figure 5B). With 460,651 4-fold symmetric particles, collected at a dose of 92 e^-^/Å^2^, the effective accumulated dose for the reconstruction is around 1.7^*^10^8^ e^-^/Å^2^ which exceeds an estimated minimum dose of 10^7^ e^-^/Å^2^ [3] by more than an order of magnitude. The spectrum of a single voxel in the protein, which contains on average 1.7 atoms of carbon, clearly shows a carbon edge (Figure 5C), giving an indication that we are approaching the range of single-atom sensitivity. An important difference to the detection of calcium at its binding site is that the carbon signal is less affected by inaccuracies in particle poses, as surrounding voxels also contain carbon.

Spectral reconstructions of worm haemoglobin were produced in the same way as for RyR1. The spectrum of the sum of all voxel values of the reconstructions similarly shows carbon, oxygen, and nitrogen K-edges, as well as L2,3-edges for chlorine, potassium, and phosphorus (Figure S6). Movie 2 shows the full series of energy-loss reconstructions.

### Estimating the resolution of the RyR1 reconstruction at the carbon edge

The resolution of reconstructed energy-loss volumes can be estimated by halfmap Fourier shell correlation (FSC) analysis. For the sum of spectral reconstructions at the carbon edge (283.2 – 287.8 eV), the FSC between the sums of reconstructed halfmaps was calculated (Figure 6A). As the halves were separated according to the gold standard during refinement, we used a correlation cut-off of 0.143, yielding a resolution of 39.8 Å.

**Figure 6.**
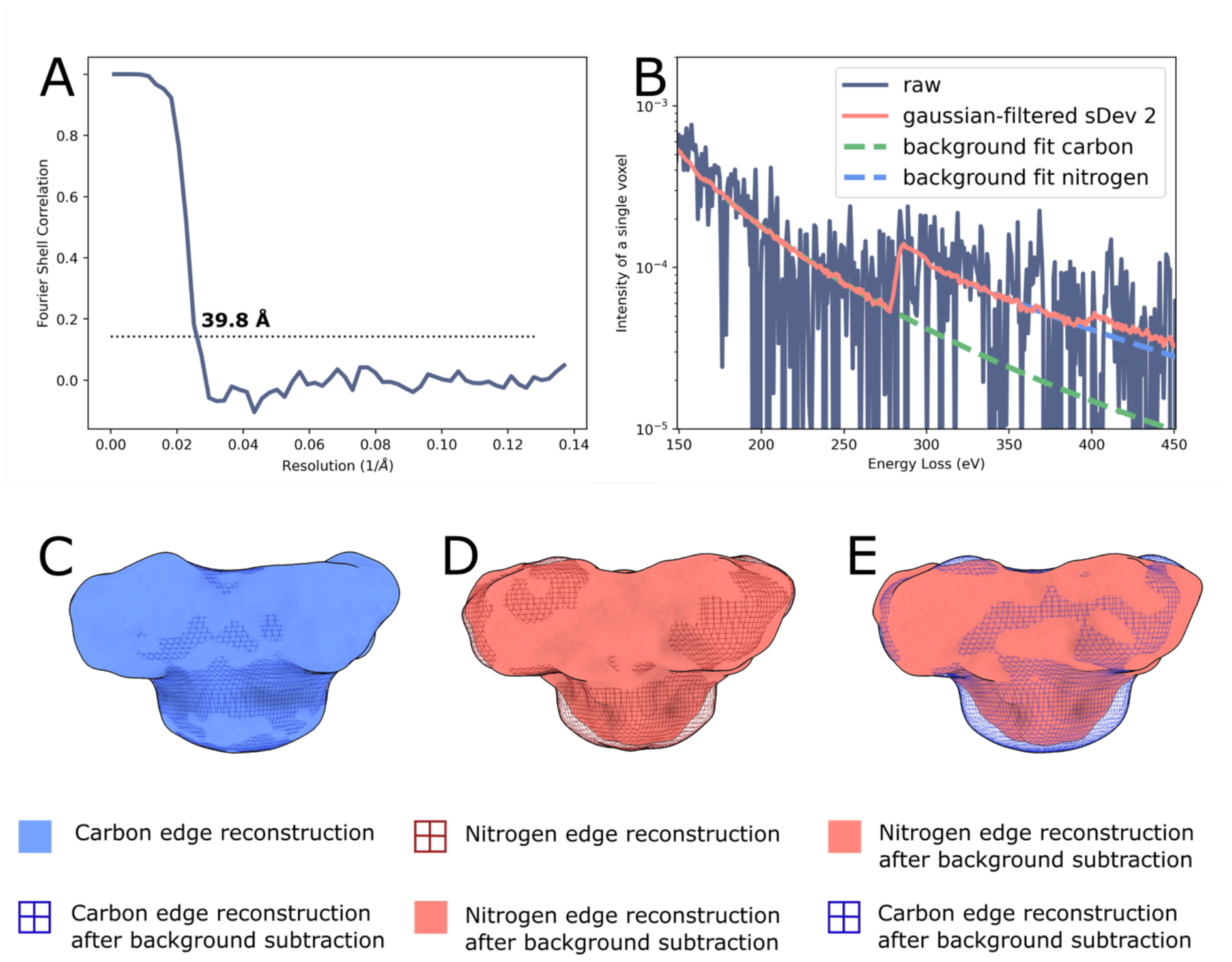
Evaluating the resolution of EELS reconstructions and the feasibility and quality of localised background subtraction. (A) The resolution of the energy-loss reconstruction at the carbon edge is estimated by reconstructing and summing halfmaps in the edge region and calculating their FSC, giving a resolution of 39.8 Å. (B) Spectrum of a single voxel of a raw reconstructions, spectrum of the same voxel after applying Gaussian filtering with a standard deviation of 2 voxels to the volumes, and background fits to the latter before the carbon and before the nitrogen edge. (C) The carbon reconstruction looks similar before and after background subtraction. (D) The nitrogen reconstruction after background subtraction shows less intensity in the region of the micelle, as the background subtraction effectively removes signal due to the long tail of the carbon edge below the nitrogen edge signal. Comparing the background-subtracted maps for carbon and nitrogen, differences in the micelle region are reflective of the lower N:C ratio in the micelle than in the protein region.

### Per-voxel Background Subtraction

To obtain an element-specific signal by EELS analysis, it is necessary to account for the background signal. In the case of carbon, the localisation of the signal with the protein in the spectral reconstructions has a clear onset at the carbon edge, showing that the elemental signal is strong enough to dominate above the background at this edge (Figure 5 C,D and Movies 1 and 2). For smaller edges, an element-specific signal can only be visualised after accounting for the signal of the decaying background in the EELS spectra. For our 4D spectral volumes, this requires per-voxel background subtraction at the relevant edge(s).In order to assess whether such a background subtraction is feasible with our data, we performed per-voxel background subtraction at the carbon and nitrogen K-edges. Individual voxel spectra show a relatively high level of noise, especially at larger energy losses (Figure 6B). To ensure sufficient signal for accurate background modelling, we applied Gaussian filtering with a standard deviation of 2 voxels to the reconstructed volumes (Figure 6B). For each voxel, we then fitted the region between 160 eV and 250 eV for carbon and between 325 eV and 375 eV for nitrogen with a power law function (Figure 6B), and subtracted the extrapolated fit from the subsequent energy bins. True elemental volumes were then generated by summing the background-subtracted values in the edge regions of carbon (283.2 – 287.8 eV) and nitrogen (403.3 – 407.9 eV). For the carbon edge, the background-subtracted volume appears very similar to the non-subtracted summed volume (Figure 6C). A larger difference is apparent for the maps at the nitrogen edge (Figure 6D). The nitrogen K-edge lies on the long tail of the carbon K-edge, so that the reconstructed volumes at the nitrogen edge also contain scattering contributions from carbon atoms. Comparing the unsubtracted and background-subtracted reconstructions, the reconstruction without background subtraction shows stronger density in the region of the micelle than the map with background subtraction, when adjusting to similar thresholds for the protein (Figure 6E). This reflects the higher carbon:nitrogen ratio of the micelle in comparison to the protein. The micelle is composed mainly of CHAPS for which the ratio is 16:1, while it is 3.6:1 for RyR1.This result demonstrates the importance and feasibility of performing background subtraction on REEL-EM volumes, in order to derive specific elemental distributions from reconstructed energy-loss volumes.

## Discussion

To fully understand macromolecular complexes at an atomic level, a technique for elemental mapping is needed. This method should ideally provide well-resolved 3D localisation, and be compatible with low-dose imaging of cryo-preserved samples while providing single-atom sensitivity.

Here, we have developed a workflow for a new technique, REEL-EM, in which EELS data are reconstructed in 3D, allowing a high total dose to be accumulated from many individual spectral images of a cryo-preserved complex, each at less than 100 e^-^/Å^2^. We present proof-of-principle data, showing that we can use poses determined from a reference refinement to reconstruct a 4D data set: a full energy-loss spectrum of 3D volumes, which correctly reproduces known distributions of abundant elements, including light elements that are inaccessible to anomalous scattering techniques. 3D per-voxel background subtraction can be performed on this 4D data set to disentangle elemental signal from background. We have compared the use of dark-field and elastic bright-field images, as well as a difference image of the two, for determination of reconstruction poses. We found the elastic bright field to give the best results, which may be related to differences in noise between the detectors used in our setup. To facilitate the collection of large data sets, we have established a workflow for automated data collection. We have further established an image processing pipeline for spectral reconstruction, using python-based tools and an adapted version of the reconstruct function of RELION, which maintains an external intensity scale between reconstructions.

The resolution of the elemental maps for the RyR1 data set can be estimated to around 4 nm on the basis of halfmap reconstructions at the carbon edge. Although this resolution does not allow neighbouring atoms to be distinguished, it is sufficient for localisation on the order of domains. In combination with a high-resolution reconstruction from TEM, even a low-resolution elemental map may be sufficient to assign unknown densities. The results presented here fall short of single-atom sensitivity, which would be desirable for most applications of the method. The cumulative dose for the RyR1 data set exceeds by an order of magnitude a previous estimate of the dose required for single-atom sensitivity by EELS [3]. However, this estimate referred to elemental detection in a single exposure. For single-particle averaging techniques, imperfect alignment accuracy increases the required cumulative dose. To distinguish between the effects of cumulative dose and the effects of alignment accuracy on sensitivity, an important observation is that, for a single voxel in the protein, containing 1-2 carbon atoms, the corresponding spectrum features a discernible carbon K-edge. This points to an important role of alignment inaccuracies in limiting the current data set.

Both image quality and processing algorithms influence the accuracy of the alignment. Image resolution in STEM depends crucially on the size of the imaging probe. Over the course of overnight data collection on our microscope, during which alignments are not re-tuned, the probe size increased significantly due to instabilities in direct alignments. This likely limits the attainable resolution for most of the images. Moreover, our reference images are formed by amplitude contrast which is less efficient than phase contrast for SPA samples, which are generally thin and composed of light elements. STEM SPA resolution can potentially be improved by phase retrieval methods [23, 24]. Practical considerations constrain the combination of these methods with EELS but hold promise for the future [25-28].Processing algorithms used in this work are standard SPA algorithms and therefore optimised for TEM phase contrast images. Adjustments for amplitude contrast images hold potential for improvements.

Regardless of possible improvements in image quality, an increase of data quantity will certainly contribute to improved elemental sensitivity. For this, an increase in data collection speed is desirable, considering that this study already required several weeks of instrument time. We therefore require faster detectors, a development that is already underway [29, 30]. Faster detectors would further allow dose fractionation and motion correction, which are currently not practical due to limitations in detector frame rate.

Reaching single-atom sensitivity would enable a deeper understanding of macromolecular complexes than is currently possible by any technique. In addition to metal ions, identification of lighter ionic species has the potential to enrich our understanding of ion channels and transporters, as well as many complexes essential to transcription and translation. Detection of sulphur or selenium substituents has the potential to guide chain tracing in poorly ordered protein regions. Mapping of lipid interactions, especially in combination with elementally labelled lipids, has the potential to enable understanding of not only stable but also transient interactions that are essential for membrane protein function.

The data presented here provide proof-of-principle for the REEL-EM method. They indicate the most important avenues for further improvements in resolution and sensitivity. With future developments, REEL-EM has the potential to enrich our understanding of the structure and function of macromolecular complexes.

## Acknowledgements

We are very grateful to Holger Stark, Max Planck Institute for Multidisciplinary Sciences, for providing infrastructure and support for the project, and to Werner Kühlbrandt and Martin Beck, Max Planck Institute of Biophysics, for support. We thank Jenn Gray (Pennsylvania State University) for collecting preliminary data and Sjors Scheres for helpful comments on the RELION code. We thank Juan Castillo, Özkan Yildiz and Tobias Koske for computing support in Frankfurt and Göttingen and the Central Electron Microscopy Facility of the Max Planck Institute of Biophysics for providing infrastructure and support for sample preparation and screening. We are grateful to David Mastronarde (UC Boulder) for the development and maintenance of the SerialEM package for automated EM data acquisition. This project has received funding from the Max Planck Society and the European Research Council (ERC) under the Horizon 2020 research and innovation programme (Grant agreement No.101116848 to B.J.M.).

## Author contributions

B.J.M. initiated and directed the project. O.P.-G., H.V.R. and B.J.M collected, processed and analysed the data. D.R. assisted in the microscopy. Y.S.C. and F.P. purified RyR1 and prepared cryo-EM samples. O.P-G. and B.J.M. prepared WH samples. D.L., P.K., M.L., and H.M. assisted with the energy filter integration and automated acquisition. O.P.-G. and B.J.M drafted the manuscript and revised it in consultation with all other authors.

## Data Availability

All maps shown in this work have been deposited to publicly accessible databanks. For the RyR1 data set, the reference reconstruction and corresponding half maps, the summed pre-carbon, carbon, nitrogen, and oxygen maps, as well as the background-subtracted carbon and nitrogen maps, and the summed carbon halfmaps were deposited to the EMDB under accession code EMD-19191. The full spectrum of energy-loss reconstructions is available at EDMOND under DOI 10.17617/3.X1R0AQ. For the WH data set, the reference reconstruction and corresponding halfmaps, and the summed pre-carbon, carbon, and oxygen maps were deposited to the EMDB under accession code EMD-19190. The full spectrum of energy-loss reconstructions is available at EDMOND under DOI 10.17617/3.AUERWM.

## Code Availability

Scripts used in the acquisition and processing of the data presented are based upon freely available software packages and are included in the supplementary materials.

## Methods

### Sample Preparation

Rabbit RyR1 was purified as described previously [31] with modifications. All steps were performed at 4 °C. In brief, 200 g of frozen rabbit skeletal muscle tissues (Pel-Freez Biologicals) were blended for 120 s in 2 x 400 mL of buffer A (20 mM Tris-maleate pH 6.8, 10% sucrose, 1 mM DTT, 1 mM EDTA, 0.2 mM PMSF, 1 mM benzamidine) and centrifuged at 3,000 g for 10 min. The supernatant was filtered through a cheesecloth and ultracentrifuged at 200,000 g for 45 min. The membrane pellet was collected, flash-frozen in liquid nitrogen, and stored at -70 °C for later use. The membrane fraction after thawing was solubilized in buffer B (50 mM HEPES pH 7.5, 1 M NaCl, 1 mM EGTA, 2 mM TCEP, 0.1 mM PMSF, 1% CHAPS, 0.2% soybean L-α-phosphatidylcholine (PC), 1:1000 of protease inhibitor cocktail (PIC; Millipore Sigma 539134)) for 1 hr. Then, an equal volume of buffer C (buffer B lacking 1 M NaCl) and His-GST-FKBP12.6 (10 mg) were added and this was stirred for another hour. The solubilized mixture was ultracentrifuged at 200,000 g for 45 min. The supernatant was mixed with 3 mL of Glutathione Sepharose 4B resin (Cytiva) and stirred for 1 hour. The resin was poured into a gravity column, washed with buffer C (50 mM HEPES pH 7.5, 0.5 M NaCl, 2 mM EGTA, 2 mM TCEP, 0.1 mM PMSF, 0.5% CHAPS, 0.1% soybean PC, 1:1000 of PIC) and eluted with buffer C plus 20 mM glutathione. The His-GST tag was removed by overnight incubation with Tobacco Etch Virus (TEV) protease (1 mg). The cleavage product was concentrated and applied to a Superose 6 Increase 10/300 GL column (Cytiva) in buffer D (20 mM HEPES-KOH pH 7.5, 250 mM KCl, 40 µM free Ca^2+^ buffered with 2 mM EGTA, 0.375% CHAPS, 0.001% DOPC, 2 mM TCEP, 0.1 mM PMSF, 1:1000 PIC). Peak fractions containing RyR1 were combined and concentrated to ∼15 mg/mL (measured by NanoDrop). RyR1-FKBP12.6 sample was mixed with calmodulin (CaM) (equilibrated in Buffer D) to a final concentration of 12 mg/mL and 100 µM, respectively. Holey carbon grids (Quantifoil Cu 300 mesh, R2/2) were prepared with fiducials by glow-discharging, adding 3 µL of 10 nm Protein A Immunogold solution and blotting from the back. The grids were left to dry overnight in a desiccator. 2.5 µL of RyR1-FKBP12.6-CaM sample were applied and blotted for 3 s (blot force of 6 to 9) using ashless blotting paper and the grids were subsequently plunge-frozen in liquid ethane using a Vitrobot Mark IV (Thermo Fisher Scientific) at 4 °C and 100% humidity.

For the worm haemoglobin sample, earth worms (Canadian Night Crawler) were purchased from a fishing supplier. Blood was extracted from the seventh segment and diluted with saline solution of 50 mM NaCl and 10 mM MgCl2. C-flat grids with a hole size of 1.2 µm and a hole spacing of 1.3 µm were used for this sample. The grids were prepared with fiducials by glow-discharging, adding 3µL of 10 nm Protein A Immunogold solution to the carbon and blotting from the back. The grids were left to dry over a minimum of one night before they were used for sample preparation. They were then again glow-discharged and 3 µL of sample were applied, before blotting and plunge-freezing in liquid ethane in a Vitrobot Mark IV, using blot force 0 and 8s blot time at 4 °C and 100% humidity.

### Spectral Image Data Collection

Data were recorded on a Titan Krios G3 with STEM capability, equipped with HAADF (Fishione) and ADF (Thermo Fisher Scientific) detectors, a CEFID energy filter [17] and a post-filter Dectris ELA hybrid pixel detector [18]. For the acquisition of spectral images, scan generation, beam-blanking and data compression were performed by the CEOS software Panta Rhei, with scanning controlled through an external scan generator (TVIPS). For acquisition of overview, alignment, and focus images, the Thermo Fisher Scientific scan generation and beam blanking were used. The experiments were performed at an acceleration voltage of 200 kV, convergence angle of 6 mrad, spot size 10, gun lens 3, 50 µm C2 aperture, and a magnification of 80 kx, corresponding to a pixel size of 3.65 Å. The collection angle for electrons entering the energy filter corresponded to 8 mrad. This was limited by the hole in the dark field detector which was inserted during acquisition, as this was smaller than the entrance aperture of the filter. To determine the dose rate, we first calibrated the average number of pixels activated per electron hit by looking at a sparse region of a spectrum, where single electron hits were clearly discernible as clusters of activated pixels. This is enabled by the linear count response of the detector to electrons up to 5×10^6^ counts/pixel/s which was not exceeded in the experiment. We then divided the total electron counts per time in an image of the beam over vacuum by this factor and the pixel area. The pixel size was calibrated by maximising the 0.5 Fourier shell correlation between an extracted bright-field reconstruction and a TEM reconstruction with a carefully calibrated voxel size of the same sample. For maximum frame rates of the detector of up to 33333 Hz, only a 66×1024 pixel area region of interest (ROI) of the detector was read out. The recorded spectra were then summed in the non-dispersive direction by Panta Rhei before saving of spectral images in hyperspy format with lossless compression. The energy filter was set to disperse 770 eV (from -25.4 eV to 745.6 eV) over the longer dimension of the detector, yielding energy bins of 0.75 eV. The bin size was later adjusted by a factor of 532/515 on the basis of the position of the oxygen edge relative to the zero-loss peak. STEM direct alignments were performed manually on an area of carbon film adjacent to the acquisition area, immediately before beginning data acquisition. Filter alignments were performed over vacuum using the automated tuning routine provided in Panta Rhei.

The acquisition was automated using SerialEM [32] for navigation and autofocus and using Panta Rhei for spectral image acquisition and saving. In SerialEM, overview images of the entire grids were acquired in TEM mode as stitched maps at low magnification. Suitable squares were then selected and recorded at an intermediate magnification in STEM mode, on the DF2 darkfield detector. Intact holes were selected as acquisition positions on the grid square maps. An automated acquisition routine was then triggered, which functions as follows: At each acquisition position, a SerialEM script moves the stage to the coordinates and the hole is further aligned on the basis of a low-dose image from the DF2 detector. The STEM autofocus routine is then run on an adjacent area of carbon and an external RPC script is triggered. SerialEM relies on the internal scan generator of the microscope (Thermo Fisher). The RPC script calls functions for spectral image acquisition and saving in Panta Rhei. This requires a switch to the external (TVIPS) scan generator which is automatically performed by Panta Rhei. Files are saved in hyperspy format. After a wait time of 12 minutes, when the current spectral image acquisition has been completed but data are still being processed and saved by Panta Rhei, the SerialEM script resumes and moves the stage to the next imaging position where the autofocus routine is again started. A wait time is set to ensure a minimum of 23 minutes between one call of the RPC script and the next to ensure that the previous file is saved before starting a new acquisition.

1142 4k x 4k x 1024 spectral images of RyR1 and 61 spectral images of WH, of which 7 included a simultaneous dark-field image, were collected with a dwell time of 30 µs per pixel at a calibrated total dose of 92 e^-^/Å^2^. The collection angle for the dark field image was 15 – 150 mrad.

### Reference reconstructions for pose determination

Using the Hyperspy framework [33] (version 1.6.4), the mean zero-loss peak (ZLP) position was calculated for every pixel in each spectral image by determining the location of maximum intensity. We found that the position was stable within the instrumental limits on energy resolution throughout each spectral image, so the mean position for each spectral image was used for the correction of the ZLP position. Elastic bright-field images were extracted as tiff files from an energy range of +-3.5 eV around the zero-loss peak centre. This encompasses 99 % of the peak. Within these micrographs, particle positions were determined using crYOLO [19] with a manually trained model. Particles were extracted and further processed in RELION-4 [20]. No motion correction, dose weighting, or CTF correction was applied. 2D classification was applied for additional data set cleaning. 460,651 particles were selected for the RyR1 data set and 16,648 particles for the WH data set. The selected particles were pre-aligned using a Class3D job with a T-value of 1 and 40 iterations, using a 60 Å – lowpass filtered initial model, generated from the data for each data set, as reference, and then subjected to 3D refinement, using a soft solvent mask. C4 symmetry was applied for RyR1 and D6 symmetry for WH. The refinement resolutions were determined by gold-standard FSC calculation in a PostProcess job.

For the spectral images for which a simultaneous dark-field image was collected, 3086 particles were picked on EBF images, generated as above. An additional set of images was produced by subtracting the dark-field images from the corresponding ADF images. The particles were extracted from all three sets of images and independently subjected to alignment by 3D classification with 40 iterations. For each map, an FSC to a map on the basis of PDB-5M3L [34, 35], generated by ChimeraX’s [36] molmap command, was calculated and resolutions were estimated from a 0.5 cutoff.

### Reconstruction of Energy-Loss Data

In a second Hyperspy script, energy-loss micrographs were extracted as tiff files from the spectral images for both data sets. For each energy bin, all particles were reextracted with the pose information from the Refine3D star file. No normalisation was applied to the particle images. For reconstruction, RELION was recompiled without line 714 in src/reconstructor.cpp, which otherwise sets the Fourier origin voxel of the reconstruction box to zero. A volume was then reconstructed for each energy, using the command relion_reconstruct with two-fold or eight-fold padding and full gridding. A single reconstruction (at 284 eV) was also made without any padding for comparison. For the maps of the pre-edge region, the carbon edge, and the oxygen edge, the volumes for 237.5 – 242.1 eV, 283.2 – 287.8 eV, and 540.4 – 545.0 eV, respectively, were summed. For display, a Gaussian filter with a standard deviation of 2 voxels was applied. For the RyR1 data set, halfmaps were reconstructed for the volumes between 283.2 and 287.8 eV by applying the halfmap option in relion_reconstruct, such that the halves were separated in the same way as for the EBF reference reconstructions. The maps were then summed for all of these volumes for each half data set and an FSC curve was calculated between the sums of the halfmaps, using relion_postprocess and a suitable mask. The mean intensities of the particles at each energy were determined from all pixels in all particle images that contributed to each reconstruction of the RyR1 data set. In Figure 4 they are compared to the mean voxel values of the resulting reconstructions.

Volumes and spectra were analysed using ChimeraX [36] and the Hyperspy library, respectively. Plots were created in python, using the matplotlib library [37]. The position of the expected calcium-binding site for RyR1 in relation to the reconstructed volumes was determined on the basis of PDB-5TAQ [22, 38], which was rigid-body-fitted into the EBF reference reconstruction. The average number of carbon atoms in a single voxel of RyR1 was determined by counting all atoms in a sphere located in a protein-dense portion of the RyR1 atomic model (PDB-5TAQ [22, 38]), dividing by the volume of the sphere, and multiplying by the volume of a voxel in the reconstruction.

Movies were produced in ChimeraX by morphing between the series of reconstructions, gaussian-filtered with a standard deviation of 7.3 Å, using one frame per reconstruction. The option “constant volume” was set to “true”. Spectrum and slider were added in the programme Adobe Premier Pro.

### Background Subtraction

For background subtraction, the spectral volumes were processed with Hyperspy. Volumes were first individually Gaussian filtered with a standard deviation of 2 voxels and then combined into a 4-dimensional EELS Object. Background fitting was performed with a power law function and the fast-fitting option disabled in the region of 186 – 273 eV for carbon and 335 – 387 eV for nitrogen. The background-subtracted volumes at each energy were then individually saved as mrc files. For the carbon map, the subtracted volumes were summed from 283.2 – 287.8 eV and for nitrogen from 403.3 – 407.9 eV. For better visualisation, Gaussian filtering was again applied to the background-subtracted nitrogen map.

### Supplementary Figures

**Figure S1.**
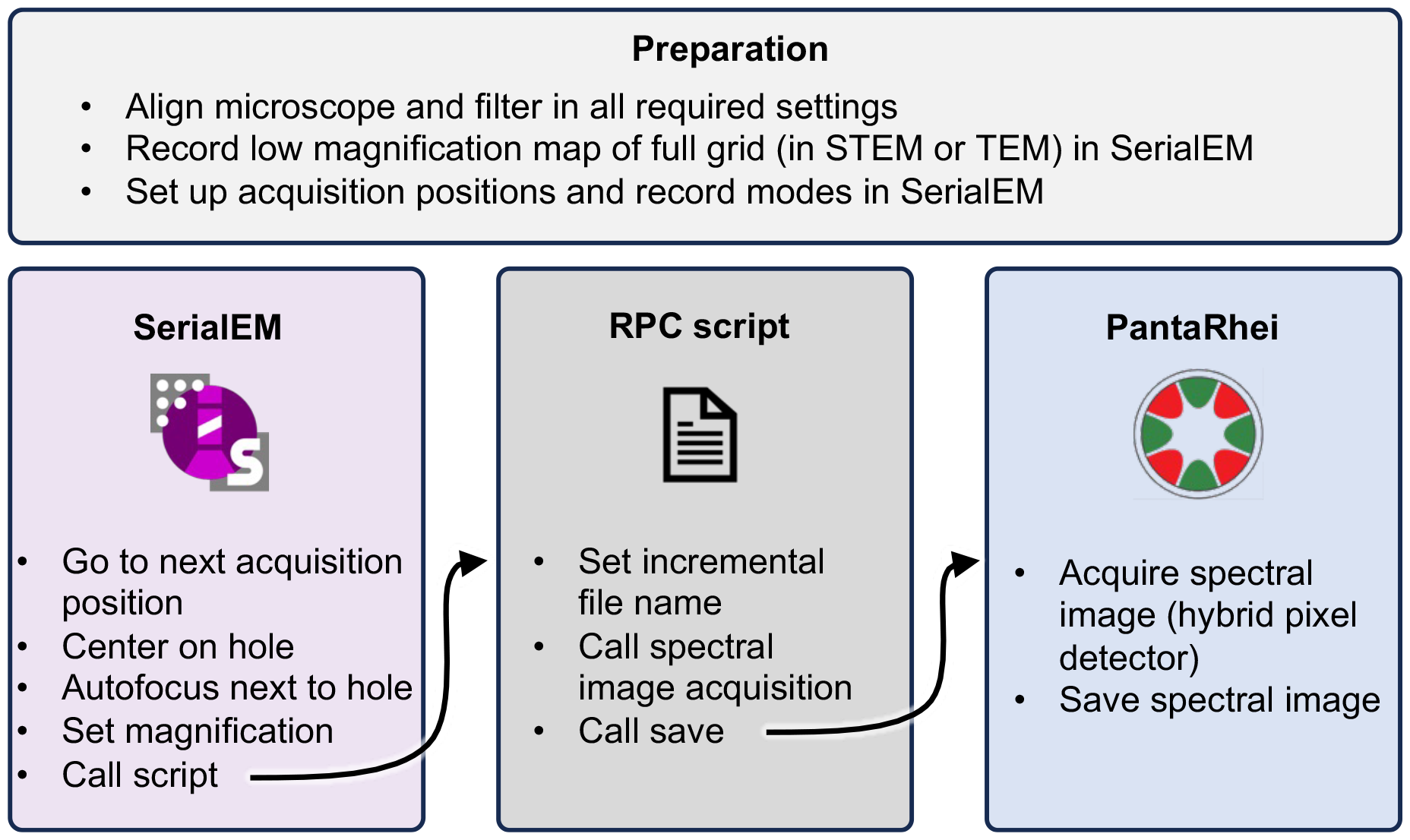
Schematic of the procedure for automated collection of spectral images. using the microscope control software SerialEM in communication with the filter software Panta Rhei via its RPC interface. After preparation steps, as indicated, automated acquisition is started. For every acquisition position, SerialEM prepares for the acquisition, then calls an RPC script which triggers Panta Rhei to acquire and save spectral images.

**Figure S2.**
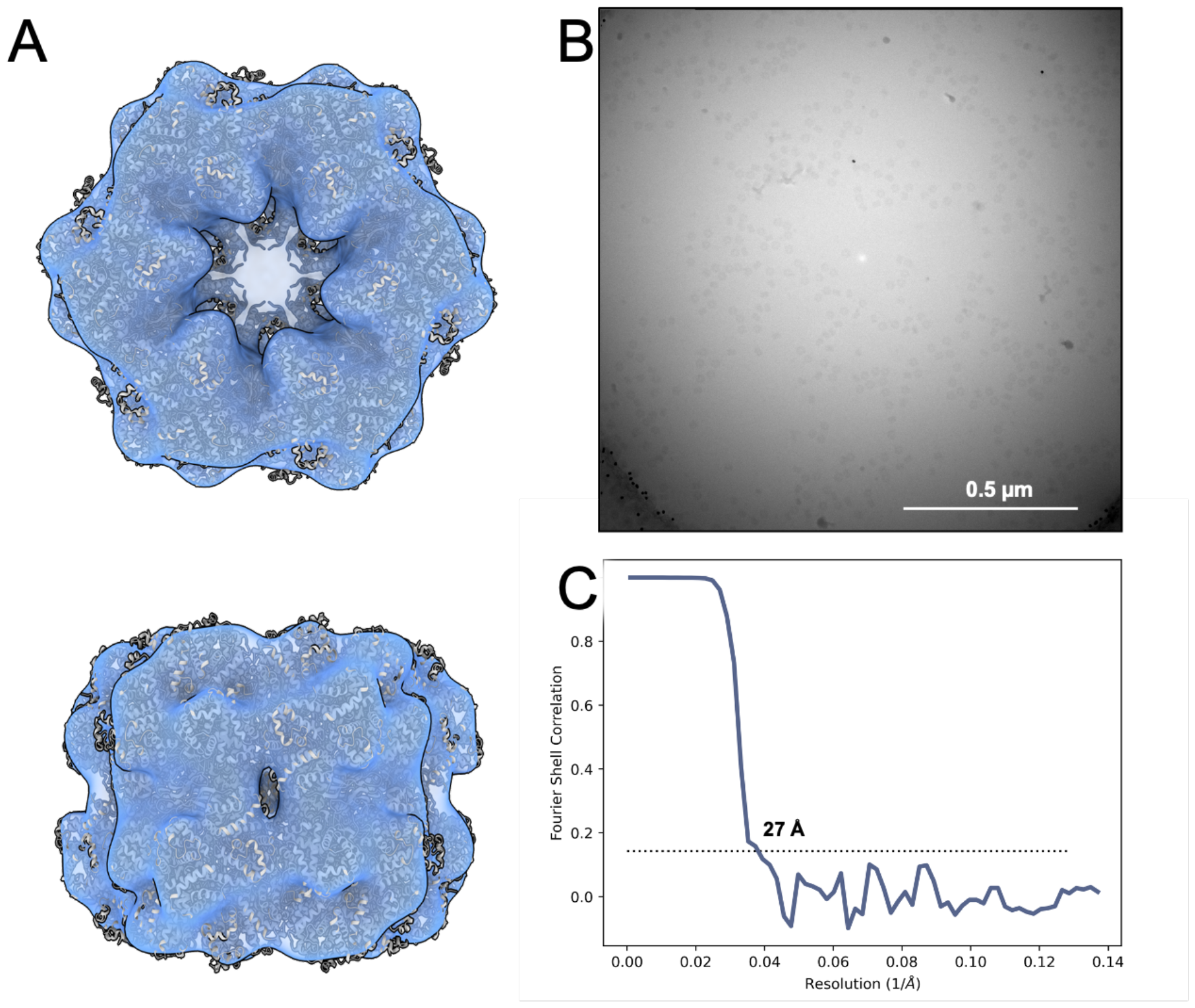
Reference reconstruction of WH. Refined reconstruction of WH from a data set of 16,648 particle images which were extracted from the zero-loss region of spectral images. The map shows a good fit – albeit at low resolution – to the published model of earthworm haemoglobin (PDB-5M3L [34, 35]) displayed as cartoon model. (B) An exemplary extracted EBF micrograph. (C) The gold standard Fourier shell correlation for this reconstruction has a 0.143 cut-off at 27 Å.

**Figure S3.**
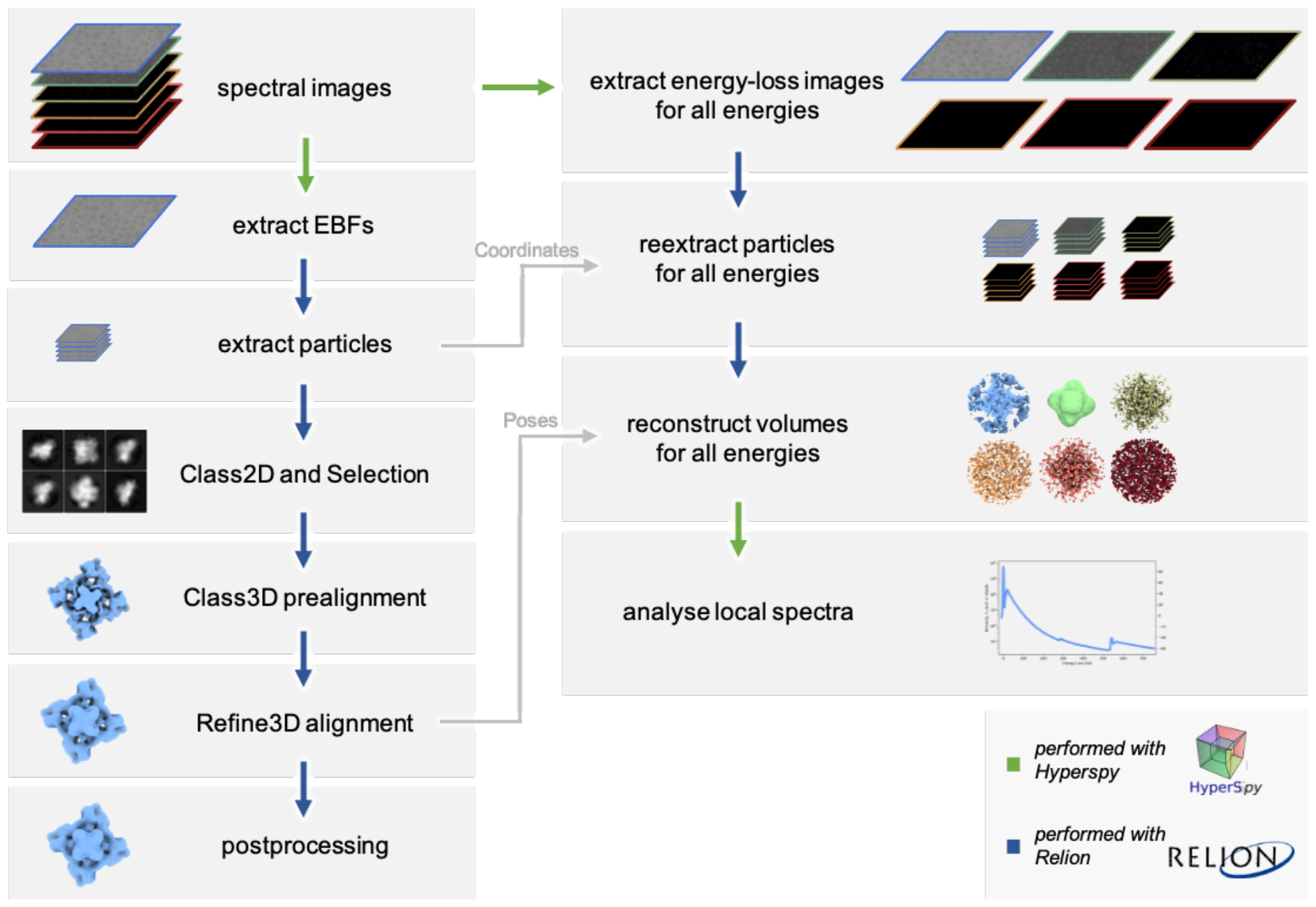
Schematic of the data processing procedure. Spectral images were processed with tools form the Hyperspy framework or the software RELION, as indicated by green or blue arrows, respectively. For the reference workflow, elastic bright field images were extracted from the spectral images. Suitable particles were picked and extracted from these, subjected to 2D classification for cleaning and then prealigned with a 3D classification job and further refined in a 3DRefine job. For resolution estimation, the results were further subjected to postprocessing. For the full spectral workflow, micrographs were extracted from the spectral images at all energies, particles were reextracted from these micrographs using the previously refined particle positions and then reconstructed for each energy, on the basis of the poses determined in the reference workflow. Spectra of specific voxels or sums of voxels were then analysed from these reconstructions in Hyperspy.

**Figure S4.**
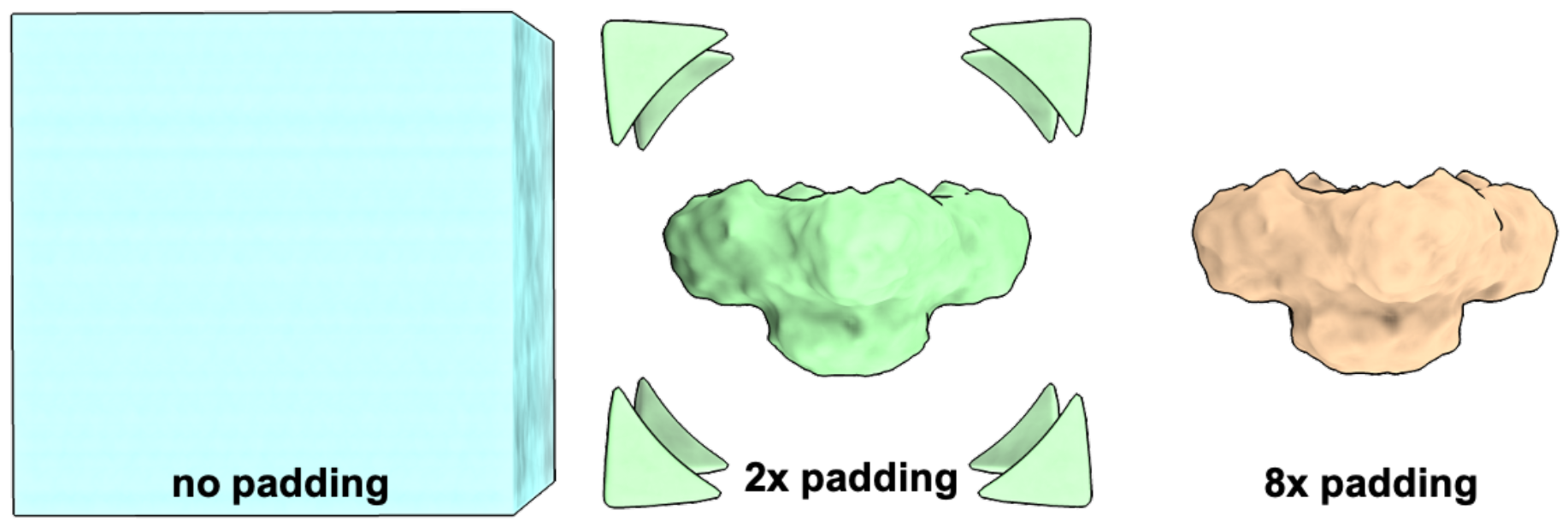
Maps reconstructed in a modified version of RELION show corner artefacts depending on padding factors. The reconstruction at 284 eV is shown for three different padding factors (Fourier volume subsampling) with Gaussian filtering with a standard deviation of 7.3 Å applied for better visualisation. No padding yields a strong artefact that dominates the reconstruction. This is reduced by padding. With a padding factor of 2, the artefact is only visible in the corners of the box, while with a padding factor of 8, the artefact does not overlap with the reconstruction at relevant thresholds.

**Figure S5.**
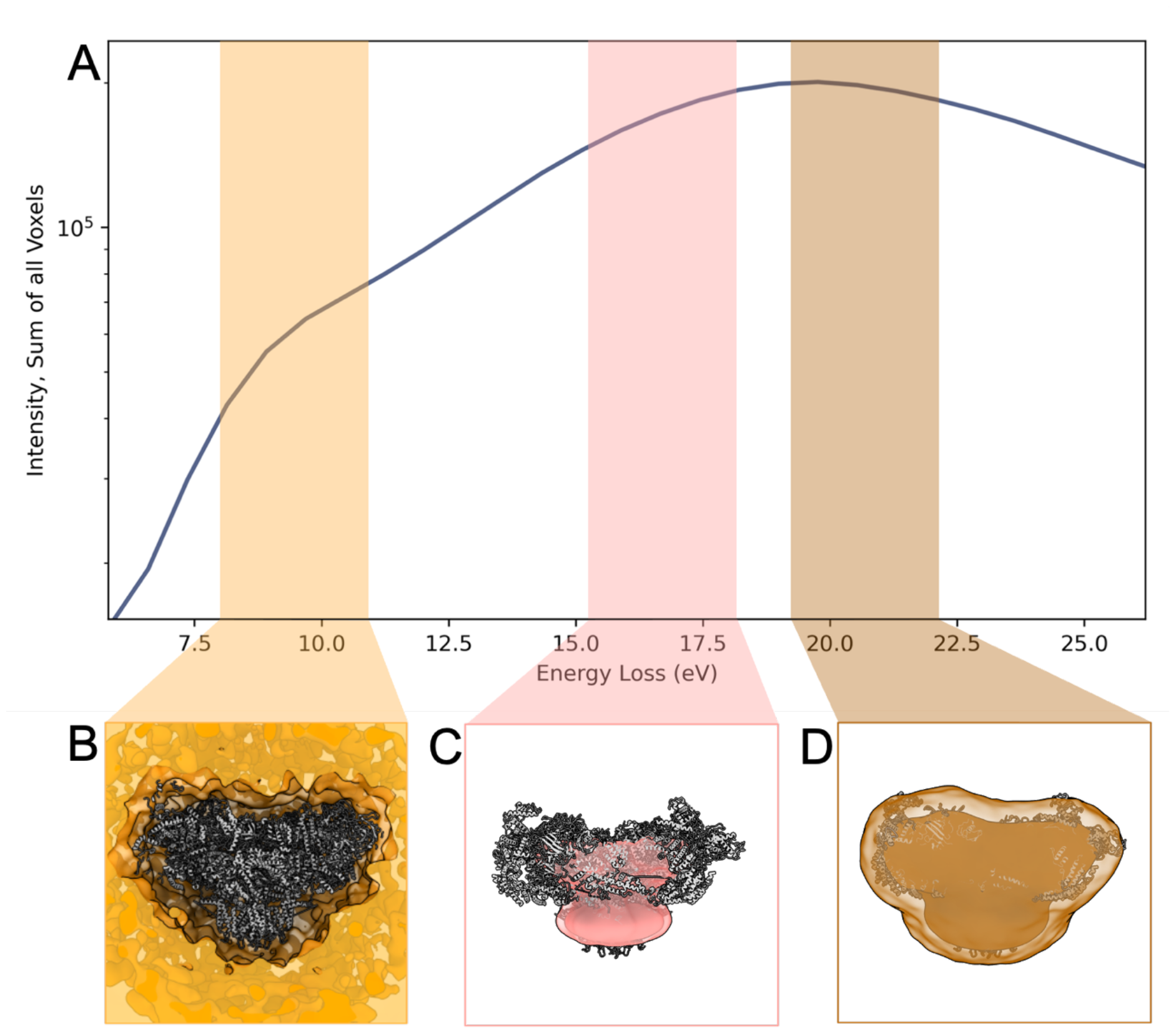
Reconstructions from the plasmon region of the energy-loss spectrum show colocalization with solvent, detergent micelle or protein locations at different energy losses. (A) Spectrum of the summed voxels of reconstructions in the plasmon region. (B-D) Summed reconstruction for the energy windows (B) 8.1 – 11.2 eV, (C) 15.1 – 18.2 eV and (D) 19.0 – 22.1, overlaid on PDB-5TAQ [22, 38], showing different localisations across the volume.

**Figure S6.**
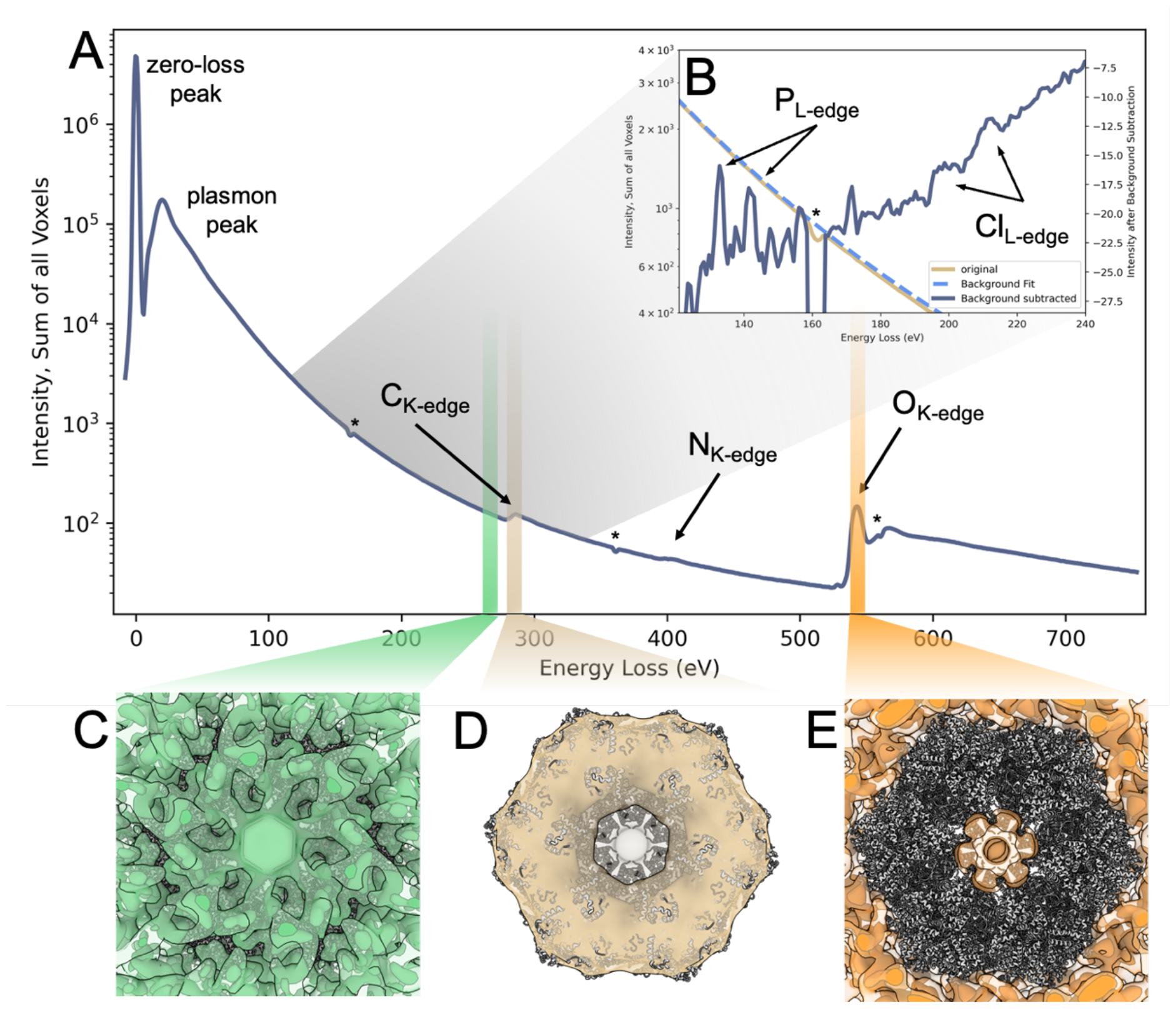
Selected spectra and reconstructions of reconstructed EELS data of WH. (A) As for RyR1, the spectrum of the sum of all intensities in the EELS reconstructions shows elemental edges for the most abundant elements: carbon, nitrogen, and oxygen. The asterisks mark regions where the spectra show dips due to double-sized pixels at the tile boundaries of the detector, which are not correctly accounted for by the detector’s flatfields. (B) For the region between 118 and 240 eV a background subtraction was performed, based on a fit of the spectrum between 121 eV and 127 eV. The subtracted spectrum shows additional edges for phosphorus and chlorine. (C-E) Gaussian-filtered reconstructions for three 4.6 eV-regions of the spectrum are shown together with PDB-5M3L [34, 35]. Before the carbon edge (C), the reconstruction shows uniform distribution of intensity between the protein and the solvent. At the carbon edge (D), the reconstruction shows colocalization with the protein and micelle, and at the oxygen edge (E), the reconstruction shows colocalization with the solvent. The reconstructions reflect the known local distributions of carbon and oxygen.

Movie 1. A morph through the full spectrum of energy loss reconstructions for RyR1. Volumes are displayed with Gaussian filtering (standard deviation 7.3 Å) for better visualisation. The thresholds for the maps were automatically adjusted during morph creation to keep the displayed volume constant.

Movie 2. A morph through the full spectrum of energy loss reconstructions for WH. Volumes are displayed with Gaussian filtering (standard deviation 7.3 Å) for better visualisation. The thresholds for the maps were automatically adjusted during morph creation to keep the displayed volume constant.

